# Physiology, functional genomics and proteomics of *Verruconatronum alginivorum gen*. *nov*., sp. nov., a first haloalkaliphilic representative of the phylum *Verrucomicrobiota from soda habitats*

**DOI:** 10.64898/2026.01.22.701064

**Authors:** Dimitry Y. Sorokin, Varada Khot, Alexander Y. Merkel, Damon Mosier, Nicole J. Bale, Michel Koenen, Marc Strous

## Abstract

Despite the successful cultivation of many microbes from rich bacterial communities inhabiting alkaline soda lakes, members of the bacterial phylum *Verrucomicrobiota* have so far been detected only through metagenomics. Here, we used alginate as a selective substrate to enrich and isolate two strains of haloalkaliphilic *Verrucomicrobiota*. The isolates share identical 16S rRNA gene sequences representing a new genus lineage, and, together with other metagenome assembled genomes, a new family within *Opitutales*. Cells of strains AB-alg1^T^ (from soda lakes) and AB-alg4 (from soda solonchak soils) are small and motile cocci forming submerged colonies in soft alginate agar. They are saccharolytic heterotrophs growing aerobically on polysaccharides (alginate, starch and inulin) and sugars (glucose, fructose, mannose, sucrose, melezitose, maltose and cellobiose). They also grow anaerobically by fermentation of alginate and D-mannose and by coupling incomplete denitrification to oxidation of alginate. Both isolates are obligately alkaliphilic and moderately salt-tolerant. The dominant membrane phospholipids include phosphatidylcholines and diphosphatidylglycerols (cardiolipins). The genome of AB-alg1^T^ features polysaccharide lyases of the PL6, 7, 15, 17, 38, and 39 families for depolymerization of alginate. Based on distinct phenotype and phylogeny, we propose classification of strains AB-alg1^T^ (JCM 35393^T^=UQM 41574^T^) and AB-alg4 as *Verruconatronum alginivorum* gen. nov., sp. nov. within a new family *Verruconatronumaceae*.

**Importance:** The presented isolates are the first isolated representatives of an environmental family of *Opitutales*, part of the core microbiome of alkaline soda lakes. These bacteria feed on polysaccharides. We present the key enzymatic machinery for the polysaccharide breakdown. These enzymes are high-pH tolerant and have potential for industry applications, for example in washing powders and biomass waste recycling.

## Introduction

Soda lakes and soda solonchak soils are unique alkaline and saline habitats characterized by stable high pH due to the presence of molar concentrations of sodium carbonate and bicarbonate, selective for obligate haloalkaliphilic prokaryotes. The prokaryotic communities of soda lakes and to a lesser extent of soda solonchak soils have been intensively studied during the last several decades in Central Asia (southern Siberia, north-eastern Mongolia and Inner Mongolia), East African Rift Valley, California and Nevada (US) and British Columbia in Canada. Culturing and molecular ecology investigations revealed taxonomically diverse microbial communities active in carbon, sulfur and nitrogen cycling present in these soda lakes (Grant and Jones, 2016; Sorokin et al., 2014; Sorokin et al., 2015; Sorokin 2017; Vavourakis et al., 2016; Vavourakis et al., 2018; Vavourakis et al., 2019; Zorz et al., 2019; Haines et al., 2023).

Our recent efforts focused on the degradation of polysaccharides in soda lakes resulted in the isolation of a range of extremely halophilic natronoarchaea capable of growth with chitin, cellulose and many other polysaccharides of neutral sugars (Sorokin et al., 2015; Sorokin et al., 2018; Sorokin et al., 2019; Elcheninov et al., 2023). However, these organisms were unable to degrade acidic polysaccharides based on uronic acids. Therefore, attempts continued to target acidic polysaccharides-utilizing bacteria in soda lakes and soda solonchak soils. Sodium hyaluronate (a linear heteropolymer of *N*-acetylglucosamine and glucuronic acid) turned out to be an extremely selective substrate for enrichment and isolation, at moderate salinity, of haloalkaliphilic bacteria belonging to the phylum *Planctomycetota*, described recently as a new genus *Natronomicrosphaera* within the family *Phycisphaeraceae* (Sorokin et al., 2025).

Here we used the acidic polysaccharide alginate, a heteropolymer of mannuronic and guluronic acids, to enrich and isolate members of the phylum *Verrucomicrobiota* which, so far, lacked any haloalkaliphilic representative in pure culture. The isolates formed a new genus and family within the order *Opitutales*. This paper describes their phylogeny, physiology, membrane lipid composition, and results of functional genome analysis.

## Materials and methods

### Inoculum and enrichment conditions

The top 1-2 cm layer of oxic sediments and bottom brines (1:3, v/v) were collected from six moderately saline soda lakes located in the south of Kulunda Steppe (Altai region, Russia; N52^°^06’/ E79^°^09’; N51^°^37’/ E79^°^50’; N51^°^39’/ E79^°^48’; N51^°^40’/ E79^°^54’-E79^°^54’) in July 2022. The salt concentration of the brines ranged from 50 to 150 g l^-1^, the pH from 10.1 to 10.6 and the carbonate alkalinity from 0.5 to 1.8 M. 5 ml of the sediment : brine suspensions from individual samples were mixed in equal proportions in a 50 ml Falcon tube, homogenized by vigorous shacking and incubated statically for 1 h to allow sedimentation of coarse heavy particles. The residual top 10 ml fraction containing fine particle suspension was then used as inoculum (5% v/v). The inoculum from 3 soda solonchak samples (collected in the same area as the lakes) was prepared by the same way: 1 g (dry weight) of each soil were mixed and homogenized in a sterile mortar to a fine powder and suspended in sterile 0.5 M NaHCO_3_ 1:5 (w/v). After vigorous shacking, heavy sand material was sedimented and the residual fine fraction was used as inoculum (5% v/v).

The enrichment medium was based on sodium carbonate/bicarbonate buffer containing 0.1 M NaHCO_3_, 0.2 M Na_2_CO_3_, 0.1 M NaCl, 1 g l^-1^ K_2_HPO_4_ and 20 mg l^-1^ yeast extract (final pH of 9.5 after autoclaving at 120 ^°^C). After sterilization, the base medium was supplemented with 1 mM magnesium sulfate, 4 mM NH_4_Cl and 1 ml each of trace metal and vitamin mix (Pfennig and Lippert, 1966; Pfennig and Trüper, 1992). 1 g l^-1^ sodium alginate from brown algae (Sigma-Aldrich) was added from a 5% (w/v) stock solution as the carbon and energy source, and cultures were incubated at 30 ^°^C on a rotary shaker at 120 rpm in screw cap bottles until visible microscopic evidence of bacterial growth (after settling of sediment particles by low-speed centrifugation).

### Growth physiology

Substrate profiling was performed in aerobic cultures at optimal pH/salt conditions and 30 ^°^C (see below) in serum bottles (5 ml/23 ml) with rubber stoppers to prevent medium evaporation, shaken at 150 rpm. Substrates were added from 10% sterile stocks (polysaccharides – autoclaved, sugars – filter-sterilized). Growth intensity was estimated by increase of OD600 in comparison to the cell-free controls, either directly in case of soluble substrates or after precipitation of insoluble polysaccharides either by gravity or by a low speed centrifugation. Salinity tolerance was examined in a sodium carbonate/bicarbonate buffer (pH of 9.5) containing 0.1-2.5 M Na^+^. The growth pH range was studied using 50 mM HEPES/50 mM potassium phosphate/0.6 M NaCl for the neutral range from 6 to 8, 0.5 M bicarbonate/1.M NaCl for the intermediate pH 8-8.5 and bicarbonate/carbonate with 0.6 M total Na^+^ for the alkaline range from pH 8.5-10.5. The actual pH was measured at the start and at the end of the experiments since it was drifting substantially from the starting set up values at the lowest and highest extremes. These incubations were performed at 30 ^°^C on a rotary shaker at 150 rpm. All other growth tests (temperature range, substrate utilization profiling, anaerobic growth) were done in carbonate medium with 0.6 M total Na^+^ and pH 9.5. Anoxic medium was prepared using a sterile Argon gas flushing-cold boiling evacuation system (3 cycles) in 23 ml serum bottles with 10 ml medium closed with butyl rubber stoppers. The medium was made anaerobic by a final addition of 0.1 mM of filter-sterilized Na_2_S. Biomass growth was monitored by increase in OD_600_. A rotary shaker (150 rpm) was used for aerobic incubations, while anaerobic growth was conducted in static incubations.

### Microscopy and chemotaxonomy

Phase contrast microscopy (Zeiss Axioplan Imaging 2 microscope, Göttingen, Germany) was applied for routine checks. Electron microscopy was used to examine flagellation and cell ultrastructural organization. For the latter, the cells were centrifuged, resuspended in 0.5 M NaCl and fixed with *p*-formaldehyde (final concentration 3%, v/v) at room temperature for 2 h, then washed again with the same NaCl solution. For whole cell imaging, the fixed cells were positively contrasted with 1% (w/v) uranyl acetate. For thin sectioning, additional fixation was done by 1% (w/w) OsO_4_. After washing in 0.5 M NaCl (pH 7), the cells were embedded in 2% agarose, dehydrated in acetone, and embedded in Araldite. Thin sections were sequentially stained with 2% uranyl acetate and 2% lead citrate for 20 min each and examined with a Jeol JEM-1400 electron microscope (Japan).

Membrane polar lipids and respiratory quinones were extracted with a modified Bligh-Dyer procedure from freeze-dried cells grown at 30 ^°^C at 0.6 M total Na^+^, pH 9.5 with Hyl until the late exponential growth phase and analysed by Ultra High Pressure Liquid Chromatography-High Resolution Mass Spectrometry (UHPLC-HRMS^n^), as described previously (Bale et al., 2021). For the core fatty acids profiling, the cells were hydrolyzed in HCl/MeOH (1.5 N) and extracted with dichloromethane and further processed, analyzed and lipids identified as described by Bale et al. (2019; Bale et al. 2021).

### Genome sequencing and phylogenomic analysis

The strains were first identified by the Sanger sequencing of their nearly complete (1,452 nt) 16S rRNA genes amplified with 3 forward and one reversed primer set. For short read DNA sequencing of strain AB-alg1^T^, genomic DNA was extracted with the FastDNA™ SPIN Kit for Soil (MP Biomedicals, United States). Shotgun library preparation was performed using the Illumina DNA Prep (M) Tagmentation kit and sequencing was performed on a NovaSeq 6000 (2 x 151 bp) system (Illumina, San Diego, CA, USA). For the long reads, high molecular weight genomic DNA was extracted and purified by the phenol-chloroform method in presence of CTAB according to the JGI protocol v.3. The further preparation of genomic libraries and Nanopore sequencing were performed as described previously (Sorokin et al. 2020). The contigs were assembled with Unicycler v.0.5.0 (Wick et al., 2017) and submitted for automatic annotation to the PGAP (Tatusova et al., 2016) in GenBank. Genome completeness and contamination were assessed using CheckM v1.2.4 (Parks et al., 2015). For placement of the AB-alg1 genome into a species tree, 120 single copy conserved bacterial markers were identified and aligned according to the Genome Taxonomy Database (Parks et al., 2018) from 703 genomes, including an outgroup of *Planctomycetota* aligned using GTDB-Tk v2.4.1 (Chaumeil et al., 2022) “identify” and “align” commands. These genomes were manually selected from the GTDB RS226 database to maximize the quality (based on completeness and contamination) of representative genomes for all class-level groups from the phylum *Verrucomicrobiota*, all order-level groups from the class *Verrucomicrobiia* and all family-level groups from the order *Opitutales*. A species tree was built using the resulting alignment with the IQ-TREE2 program v2.2.0.3 (Minh et al., 2020) with fast model selection via ModelFinder (Kalyaanamoorthy et al., 2017) and ultrafast bootstrap approximation with 1000 bootstraps (Minh et al., 2013). Relative evolutionary divergence (RED) value were calculated using GTDB-Tk v2.4.1 “de_novo” workflow (--gamma --prot_model LG --outgroup_taxon p Spirochaetota) and PhyloRank (https://github.com/dparks1134/PhyloRank). Percentage of conserved proteins (POCP) was calculated using POCP-nf v2.3.6 (Hölzer M., 2024). A search among high-throughput sequencing data for 16S rRNA gene regions was performed by IMNGS (Lagkouvardos et al., 2016) using 99% similarity threshold and 200 bp as a minimum size threshold. The presence of MAGs in public shotgun metagenome datasets was analyzed by Sandpiper 1.0.1 (Woodcroft et al., 2025) using f_T3Sed10-336 as query for search by GTDB RS226 taxonomy.

### Ancestral reconstruction analyses

Genes were predicted and annotated from the same genomes used for the phylogenetic placement of AB-alg1using MetaERG v2.5 (Dong & Strous, 2019). Seventy-four representative genomes were subsampled from the species tree. This included all the genomes, with a completeness ≥ 90% and contamination ≤ 5%, from the *T3Sed10-336* family, 3 representatives from every other family in the *Opitutales* order, 2 representatives every other order from the *Verrucomicrobiae* class, 1 representative from every other class in *Verrucomicrobiota* phylum and 5 representatives from the *Planctomycetota* phylum as an outgroup (**Table S1**). A multiple sequence alignment was created for these genomes based on 120 single copy conserved bacterial markers according to the Genome Taxonomy Database (Parks et al., 2018) from 74 genomes using GTDB-Tk v2.4.0 (Chaumeil et al., 2022) “identify” and “align” commands. A species tree was built using the resulting alignment with the IQ-TREE2 program v2.2.0.3 (Minh et al., 2020) with fast model selection via ModelFinder (Kalyaanamoorthy et al., 2017) and ultrafast bootstrap approximation with 1000 bootstraps (Minh et al., 2013) as well as approximate likelihood-ratio test for branches (Anisimova and Gascuel, 2006). Orthogroups were formed using genes from these 74 genomes using Orthofinder v3.1 (Emms et al. 2025) and aligned within Orthofinder using famsa v2.4.1 (Gudyś et al. 2025). Gene trees were then constructed for orthogroups containing genes from AB-alg1 identified to participate in the alginate metabolism using IQ-TREE2 v2.2.0.3 (Minh et al., 2020). First, a guide tree was created using the model “LG+F+G”, followed by a posterior mean site frequency (PMSF) model with “LG+C20+G” and 5000 ultrafast bootstraps (Wang et al., 2018). Gene trees were reconciled with the species tree using AleRax v1.4 (Morel et al., 2024) with either 100 or 500 gene tree samples. For further analyses, only genes with a >= 70% probability were considered present at ancestral nodes and transfer events had to be seen in at least 50 bootstraps. Reconciled gene trees were visualized using ThirdKind (Penel et al., 2022) and the ggtree package in R (Yu et al., 2017).

### Functional genomic and proteomics analyses

Polysaccharide lyase families which include characterized endo/exo-alginate hydrolyzing activities documented in the CAZy database (Drula et al., 2022; Lombard et al., 2014) were identified using the dbCAN3 portal scan of the translated proteome (Zheng et al., 2023). The identified proteins were manually checked for the presence of signal peptides using the Signal 6.0 server (Teufel et al., 2022). Transmembrane sequences were predicted using TMHMM 2.0 (Krogh et al., 2001). Similar and experimentally validated proteins were identified using Blast against the UniProt database (The Uniprot Consortium. 2025). Domain architecture was inspected with the HMMER 3.4 portal (Potter et al., 2018) and InterPro 106.0 (Blum et al., 2024). Inference of the metabolic pathway for uronate hydrolysis was done by UniProt Blast search (Altschul et al., 1997) using model proteins from an alginate-metabolizing *Flavobacterium* sp. (Nishiyama et al., 2021) and *Vibrio pelagicus* (He et al., 2022). Nitric oxide reductase was identified using custom HMM models (Murali et al., 2024).

Proteomics was performed for strain AB-alg1^T^ grown on alginate, mannose and sucrose. Cells were harvested for proteomics after 3 consecutive transfers on each of these substrates at mid-exponential growth phase. The detailed workflow protocol was as described previously (Sorokin et al., 2020). Briefly, the freeze-dried cells were lyzed in an SDS/DTT buffer at 95 ^°^C, the total proteins were separated on a Microcon spin filter and hydrolyzed by trypsin. Peptides were desalted with C18 ZipTips (Millipore Sigma) according to manufacturer’s protocol and submitted to LC-MS/MS. Data were collected using an Easy-nLC 1200 system coupled to a QExactive Plus (ThermoFisher Scientific). Approximately 1.5µg of each sample was injected onto a 300μm x 5mm PepMap Neo Trap Cartridge (C18, 5μm particle size, 100Å pore size, ThermoFisher Scientific) and then separated by reverse phase chromatography using an EASY-Spray HPLC analytical column (75μm x 75cm, C18, 2μm particle size, 100Å pore size, ThermoFisher Scientific) at a flow rate of 225 nL/min at 40 °C. Mobile Phase A consisted of 0.1% (v/v) formic acid in water, while Mobile Phase B consisted of 0.1% (v/v) formic acid in 80:20 acetonitrile:water (v/v). Separation was carried using a multistep 140min gradient from 2-31% B for 102min; 31-50% B for 18min, followed by a wash step at fixed 99% B for 20min. Mass spec data were collected in positive mode at 1800V using a data-dependent acquisition (DDA) routine. Full MS scans were performed in the Orbitrap at 70,000 resolution (at m/z 200) for the 380-1600 m/z range, with a maximum injection time of 200ms and AGC target set at 3e6. The Top15 precursors charged 2-8+ were selectively isolated (quadrupole isolation window of 1.2 Th) for fragmentation through HCD at a normalized collision energy of 25. MS2 scans were performed in the Orbitrap at 17,500 resolution (at m/z 200) at the 200-2000 m/z range, with maximum injection time of 100ms and AGC target set at 5e4. Dynamic exclusion window was set to ignore ions within 20s and 10ppm tolerance. Spectrum Peptide Matching was performed using FragPipe (Kong et al., 2017; Yu et al., 2020aYu et al., 2020b; Teo et al., 2020), with the predicted protein sequences of strain AB-alg1^T^ as the database. The raw data were deposited in the proteomic archive PRIDE under accession number PXD071159.

## Results and Discussion

### Isolation of pure cultures

Both soda lake and soda soil primary alginate enrichments showed growth (in turbidity and cell density increase) within 10 days and were dominated by small motile cocci. They were first passed two times at 1:100 dilution to ensure growth reproducibility and to obtain sediment-free cultures. That had proven to be difficult because of a development of flagellated protozoa rapidly decreasing the bacterial cell density. The protozoa were successfully eliminated by serial dilutions up to (10^-10^) in the same medium. However, both enrichments still contained a rod-shaped phenotype and surface plating resulted in colony formation only from the latter. Therefore, considering our previous experience with the hyaluronate-utilizing *Natronomicrococcus* from the same habitats, attempts were made to achieve colonial growth of the dominant micrococci by mixing the culture into soft agar. No colonies formed in 1 % agar, while use of 0.6-0.8 % agar led to formation of separate colonies of the target cocci from both enrichments after inoculation with up to 8-fold diluted liquid cultures. Transfer of a colony into liquid medium with alginate resulted in vigorous growth of morphologically uniform motile micrococci and the purity of two cultures was confirmed by the nearly complete 16S rRNA gene Sanger sequencing: the lake isolate was designated as AB-alg1^T^ and the soil culture – as AB-alg4. As their 16S rRNA gene sequences were identical, further work was mostly done with the type strain AB-alg1.

### Phylogenetic analysis, classification and environmental distribution

Preliminary identification based on the 16S rRNA gene sequence analysis placed the alginate isolates into a new lineage within the phylum *Verrucomicrobiota* which, so far, did not have any cultivated haloalkaliphilic representatives from soda habitats. The closest relatives among the cultivated members in this phylum were from the order *Opitutales*, with the percentage of 16S rRNA gene identity below 90 %.

DNA was extracted from the AB-alg1^T^ cells grown with alginate and after short read and long read sequencing, the genome was assembled into a single linear contig. The genome size was 6.01 Mb with a GC content of 59.4 % and completeness and contamination of 99.32 and 1.69 %, respectively (Parks et al., 2015). To clarify the phylogenetic position of AB-alg1^T^ strain we made a phylogenomic reconstruction based on 120 single copy marker genes (bac120) according to Parks et al. (2018), but with a much larger alignment length (23,562 aa) and using maximum-likelihood instead of approximate maximum-likelihood. This reconstruction confirmed that this bacterium forms a deeply-branching lineage within the *Opitutales*, which, together with several metagenome-assembled-genomes (MAGs) from soda lakes, forms a separate family, sister to the type family *Opitutaceae* (**Fig. 1**). This phylogenetic lineage is designated in the GTDB release RS226 as T3Sed10-336 (the abbreviation originated from the “sediments of Tanatar-3 soda lake in Kulunda Steppe). and has 100% bootstrap support in both our and the GTDB RS226 phylogenetic reconstructions. This lineage has the relative evolutionary divergence (RED) value 0.742, which is slightly below than the median RED values for bacterial families in GTDB RS226 (0.762). AB-alg1^T^ strain represents the separate genus-level lineages, in addition to which the T3Sed10-336 cluster also includes several genus-level lineages of MAGs from moderately saline soda lakes in the same area [g__PXDC01, g SLOW01 and g T3Sed10-336 (Vavourakis et al., 2018)] and from a similar type of soda lakes in British Columbia (Canada) [g JAIUMY01 (Zorz et al., 2019)]. The AB-alg1^T^ strain and the MAG closest to it (GCA_035302665.1) have only 39.8% POCP. In addition, their joining node has a RED value of 0.869, which is significantly lower than the median RED values for bacterial genera in GTDB RS226 (0.92). Thus, the AB-alg1^T^ strain represents a standalone genus. 16S rRNA based analysis using the IMNGS service showed that phylotypes related to AB-alg1 (>99% identity of the V4 region of the 16S rRNA gene) are found only as single or a few sequences in samples of two saline soda lakes: Mono Lake (USA, PRJNA387610) and Lake Gahai (China, PRJNA294836). At the same time, metagenome-based analysis using the Sandpiper resource showed that phylotypes belonging to T3Sed10-336 family-level lineage are found in ∼800 metagenomes and may represent up to ∼30% of microbial community in photobioreactors inoculated with microbial mats from Canadian alkaline soda lakes (Ataeian et al., 2022).

**Fig. 1.**
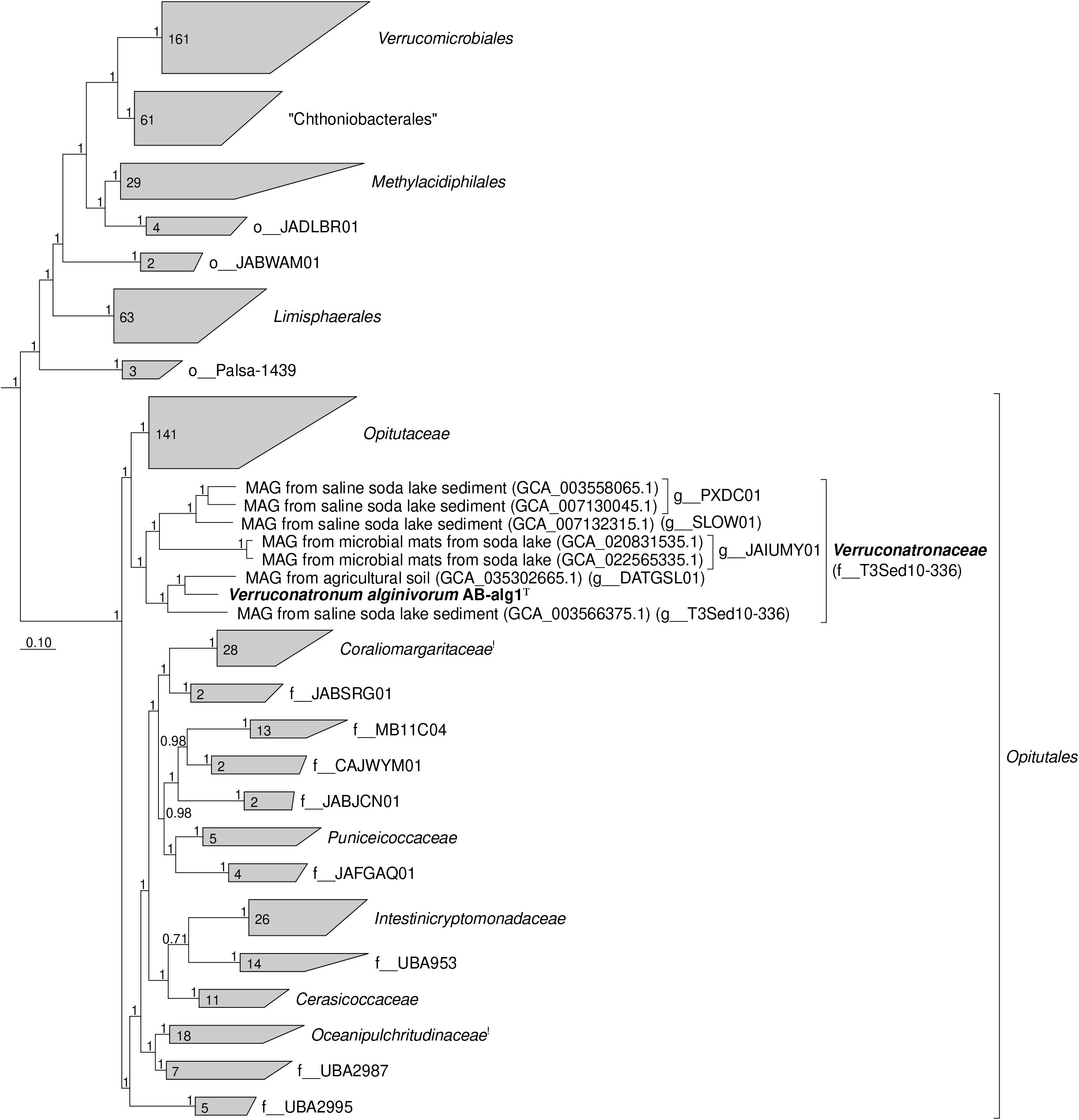
Phylogenetic placement of strain AB-alg1^T^ within the *Verrucomicrobiia* based on phylogenetic analysis of concatenated partial amino acid sequences of 120 bacterial conservative proteins (Parks et al., 2018; the length of the alignment is 23,562 aa) by maximum likelihood inference. Bootstrap values are shown at the nodes (1 = 100%). Bar, 0.10 changes per position. Taxonomic designations are given according to the GTDB RS226. Clades with alkaliphilic representatives are indicated with an exclamation mark after the taxon name. All class-level groups from the phylum *Verrucomicrobiota* were included in theanalysis (not shown). Sequences of representatives of the phylum *Planctomycetota* were used for tree rooting.

### Cell morphology and chemotaxonomy

Both isolates formed hard white colonies submerged in soft agar (**Fig. 2 a**). In liquid cultures, the cells grew mostly in homogenous suspension as small mono- or diplococci and were motile with 1-2 flagella. (**Fig. 2 b-c**). A major problem with the isolates was their rapid cell lysis in liquid cultures reaching stationary phase and culture storage at 4 ^°^C. The reason for this became obvious after electron microscopy of thin sections which showed a typical Gram-negative cell ultrastructure with a loose cell wall (**Fig. 2 d**). Storage of exponentially growing cultures with 15 % glycerol (v/v) at -70 ^°^C allowed for a long-term preservation of the isolates.

**Fig. 2.**
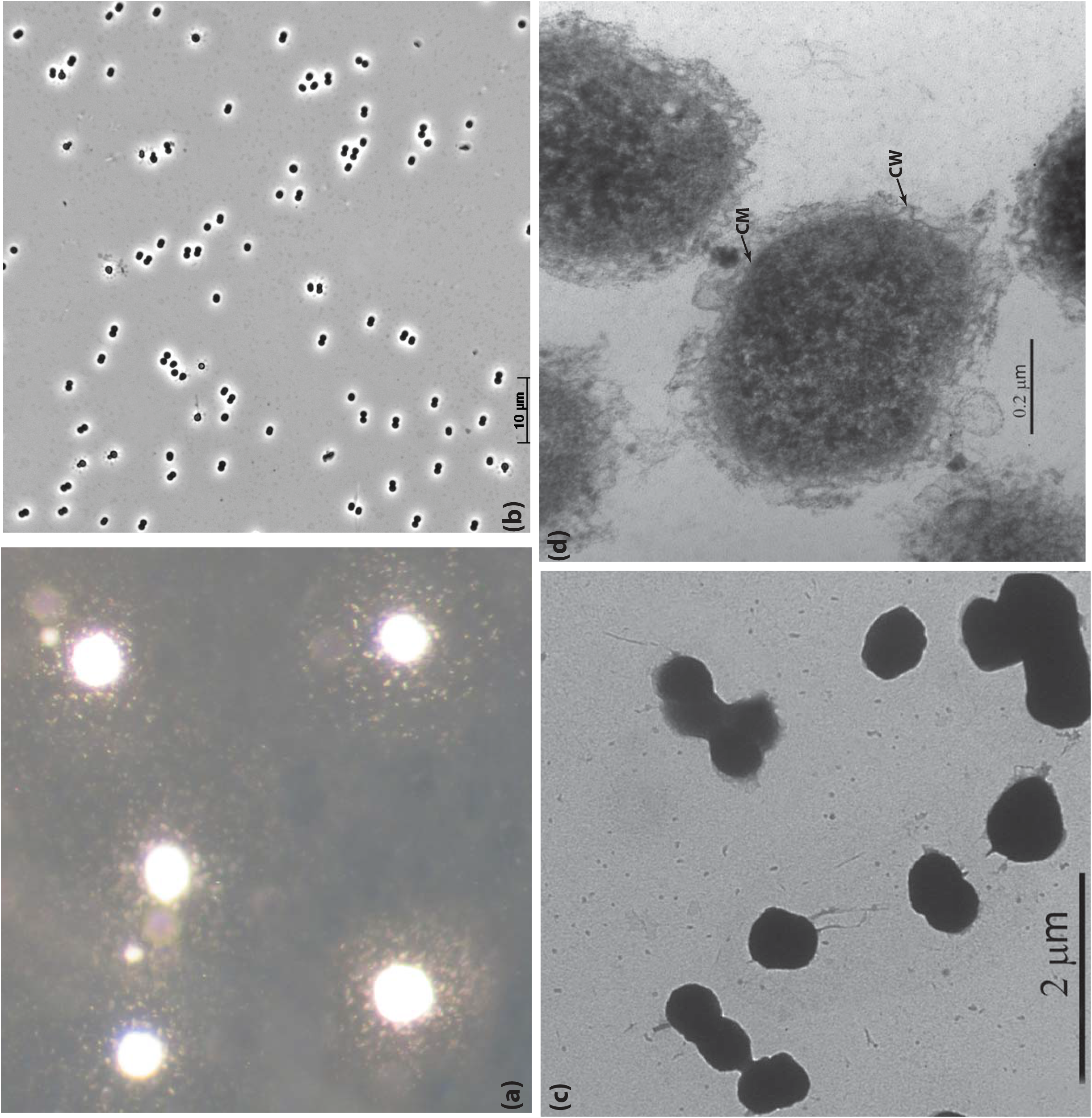
Colony (inside soft agar) (**a**), and cell morphology (**b-d**) of strain AB-alg1^T^ grown with alginate at 0.6 M total Na^+^, pH 9.5 and 30^°^C. (**b)**, phase contrast microphotograph; (**c)**, transmission electron microscopy microphotograph; (**d**), thin section electron microscopy microphotograph showing Gram-negative type of cell ultrastructure with an extended periplasm. **CW**, cell wall; **CPM** – cytoplasmic membrane.

The major respiratory menaquinones detected were fully unsaturated MK-7:7 and its 2,3-epoxy-modification, which was identified based on the mass and similarity of its mass spectrum to published spectra (Gentili et al., 2014). It is not clear, however, whether the epoxy-modification was an artifact of the extraction protocol or a natural component of the respiratory MK pool. The identified membrane phospholipids included phosphotidylcholines (PCs) and diphosphatidylglycerols (DPGs, also known as cardiolipins), with smaller amounts of phosphatidylethanolamines (PEs), lyso-PCs (which miss one fatty acid), phosphatidylglycerols (PGs) and acyl-PGs. The dominant polar lipid fatty acids (in order of abundance) were *anteiso-*C15:0, C18:1ω9, C18:0 and C16:0. Among those, the *anteiso-*C15:0 and C16:0 are also among the dominant fatty acids in other *Opitutaceae* genera, while the C18 fatty acids are absent in *Opitutaceae* but occur in other families of the *Opitutales* order **(Table 1** and **Table S2**).

**Table 1.** Comparative properties of strain AB-alg1^T^ with the type species of the related genera in the family *Opitutaceae*.

| Property | AB-*alg1*<sup>T</sup> | *Opitutus terrae*<sup>a</sup> | *Rariglobus hedericola*<sup>b</sup> | *Cefalotococcus primus*<sup>c</sup> | *Pelagococcus mobilis*<sup>d</sup> | *Ereboglobus luteus*<sup>e</sup> | *Geminisphaera colitermitum*<sup>f</sup> | *Nibricoccus aquaticus*<sup>g</sup> |
| --- | --- | --- | --- | --- | --- | --- | --- | --- |
| Colonies | white, hard* | white, granular* | white, circular | cream | white | yellow lenses | white | yellow |
| Cells | motile cocci | motile cocci | motile cocci | nonmotile cocci | motile cocci | motile cocci | motile diplococci | nonmotile cocci |
| Relation to O<sub>2</sub> | facultative anaerobe: fermentation<br>NO<sub>2</sub>>N<sub>2</sub>O | obligate anaerobe: fermentation<br>NO<sub>3</sub>>NO<sub>2</sub> | obligate aerobe | obligate aerobe | obligate aerobe | facultative anaerobe: fermentation | obligate aerobe | obligate aerobe |
| Growth substrates: polysaccharides | **alginate**, starch, inulin | starch, xylan, pectin | nd | nd | nd | pectin, | starch | dextrin |
| Growth substrates: sugars | glucose, fructose, sucrose, mannose, melezitose, fucose, maltose, cellobiose | glucose, fructose, sucrose, mannose, galactose, lactose, maltose, arabinose, cellobiose, melibiose, galacturonic acid | glucose, fructose, glucuronamide | glucose, fructose, mannose, sucrose, melibiose, maltose, cellobiose, lactose, raffinose, turanose, trehalose, | nd (only acid production) | glucose, fructose, galactose, lactose, maltose, arabinose, cellobiose, ribose, melibiose, trehalose, galacturonic acid | glucose, galactose, maltose, cellobiose | glucose, mannose, melibiose, rhamnose, maltose, cellobiose, galactose, gentibiose, lactose, lactulose, arabinose, raffinose, fucose, glucuronic acid, galacturonic acid |
| pH range (opt.) | 8.2-10.3 (9.5)** | 5.5-9 (7.5-8.0) | nr (neutrophile) | 6.9-7.7 (7.0) | 6.0-9.0 (7.5-8.0) | 6.0-9.0 (7.0) | 5.5-7.5 (7.0) | 5.5-9.0 (7.0) |
| Salinity range (opt.), Na<sup>+</sup> (M) | 0.2-1.8 (0.4-0.6) | nd | 0-0.1 (0) | 0.1-0.25 (0.15) | nd | nd | 0-0.25 (0) | 0-0.1 (0) |
| N-source (apart from ammonium) | urea, nitrate (growth, genomic) | urea (genomic) | urea, nitrate (genomic) | urea (genomic) | urea (urease test; genomic) | - | urea (genomic) | urea, nitrate (urease test, genomic) |
| Nitrogenase | - | - | - | - | - | - | +(growth, genomic) | +(activity, genomic) |
| Max. temperature (°C) | 40 | 37 | 30 | 37 | 37 | 37 | 35 | 37 |
| Catalase/oxidase tests | +/+ | -/- | -/- | -/- | -/+ | -/- | -/- | -/- |
| Intact membrane polar lipids | PC, DPG, PE, PG | nd | PE, PG, DPG | nd | nd | nd | nd | PE, PG, DPG |
| Predominant polar lipid fatty acids | *ai*C15:0, C18:0, C16:0 | nd | C14:0, C16:0, *i*C14:0, *ai*C15:0 C16:1ω5 | *ai*C15:0, *i*C14:0, C16:0, C16:1ω5 | *ai*C15:0, C16:0, C16:1ω7 | nd | *ai*C15:0, *i*C15:0 | *i*C14:0, *ai*C15:0, C16:0, *i*C16:0 |
| Respiratory lipoquinones | MK-7:7 | nd | MK-7 | nd | MK-7 | nd | nd | MK-7 |
| Genome size (Mbp) | 6.0 | 6.0 | 4.1 | 2.4 | 7.5 | 4.2 | 5.2 | 4.7 |
| G + C (% , whole genome) | 59.3 | 65.5 | 60.6 | 63.0 | 53.5 | 59.7 | 60.5 | 62.0 |
| Habitat | soda lakes and soda solonchak soils | paddy soil | freshwater | ants gut | marine | cockroach hindgut | termite gut | freshwater |

\*submerged in soft agar, \*\*actual measured values.
Lipids: PC phosphatidylcholine; PE phosphatidylethanolamine, PG phosphatidylglycerol, DPG diphosphatidylglycerol.
<sup>a</sup> Chin et al., 2001; <sup>b</sup> Pitt et al., 2020; <sup>c</sup> Lin et al., 2016; <sup>d</sup> Yoon et al., 2007; <sup>e</sup> Tegtmeier et al., 2018; <sup>f</sup> Wertz et al., 2012; <sup>g</sup> Baek et al., 2019.

### Growth physiology

Substrate profiling of strain AB-alg1^T^ and AB-alg4 demonstrated that the bacteria have an organoheterotrophic, saccharolytic metabolism with a relatively narrow spectrum of carbohydrates supporting growth. From the tested polysaccharides, the most active growth was observed on alginate. Weaker growth (with highly aggregated cells) – on soluble starch (alpha-glucan) and inulin (beta-fructan) was observed. Mannose and fucose supported excellent growth, while growth on glucose, fructose, sucrose, melizitose, cellobiose, and maltose was less vigorous. The type strain (in contrast to AB-alg4) also grew with melibiose and raffinose, but the cells showed signs of osmotic stress (being swollen and lysing rapidly). During growth on all these substrates cells were highly aggregated. The following polysaccharides tested negative as substrates: agarose, gellan, heparin sulfate, chondtroitin sulfate, fucoidan, hyaluronan, pectin (citrus or apple), polygalacturonate, rhamnogalacturonan, starch, dextran, beech xylan, xyloglucan, arabinan, galactan, amorphous chitin and cellulose, beta-mannan. The monosugars, alcohols and organic acids tested but not utilized included arabinose, galactose, rhamnose, raffinose, trehalose, lyxose, xylose, lactose, ribose, glucosamine, *N*-acetylglucosamine, mannitol, acetate, propionate, pyruvate, lactate, succinate, malate and fumarate. Anaerobic fermentative growth occurred with mannose and alginate. Yeast extract and peptones from meat or casein neither supported growth as sole substrates nor showed stimulatory effect on growth with alginate.

Anaerobic respiration was tested with 4 mM nitrite as the electron acceptor and alginate or mannose as carbon and energy sources in comparison with the fermentative growth without nitrite. The results were positive only in case of alginate: nitrite was completely consumed with the final growth yield doubled compared to fermentation. The product was nitrous oxide (N_2_O). Dissimilatory ammonification was not observed.

Salinity (in the form of sodium carbonates) and pH (at 0.6 M total Na^+^) profiling (**Fig. 3**) characterized the new isolate as a moderately salt-tolerant obligate alkaliphile. The salinity range for aerobic growth on alginate was 0.3-1.8 M total Na^+^ (optimum at 0.4-0.6 M). The pH range that supported growth was 8.2 to 10.3 (optimum at 9.5). The optimum growth temperature at pH 9.5 and 0.6 M total Na^+^ was 30 ^°^C with no growth above 40 ^°^C. A phenotypic comparison of AB-alg1^T^ and other *Opitutaceae* genera is presented in **Table 2**. What set the new isolates apart was their obligate haloalkaliphilic nature, which was unique within both the *Opitutales* order and the *Verrucomicrobiota* phylum as a whole. Note that previously reported growth of the *Opitutales* genera *Congregicoccus, Horticoccus* and *Oleiharenicola* at pH 11 remains unsupported by actual pH measurements, while use of an unbuffered medium compromised conclusions (Chung et al., 2022; Islam et al., 2025; Rochman et al., 2018). Another feature of the new isolates, distinguishing them from the majority of cultured members of *Verrucomicrobiota*, was their ability to use alginate as a growth substrate. Only hydrolysis of alginate, but not growth, has been reported previously in *Oceanipultrichrido coccoides* (Feng et al., 2020).

**Table 2.** Key functional proteins encoded in the genome of strain AB-alg1^T^ potentially involved in alginate metabolism. CBM – carbohydrate binding module. In **bold**: proteins expressed during growth on alginate, mannose, or sucrose. Gray shading indicates proteins upregulated during growth on alginate relative to growth with mannose or sucrose. The complete data is provided in **Table S3**.

| Locus tag | CAZy PL family | Protein | Putative function | Signal peptide |
| --- | --- | --- | --- | --- |
| Alginate depolymerisation |  |  |  |  |
| 000055 | PL39+CBM32/35/96 | Alginate lyase | Endo-alginate lyase | Sec/SPI-SPII |
| 000060 | PL15 |  | Exo-acting alginate lyase | Sec/SPI-SPII |
| 000395 | PL15 |  | Exo-acting alginate lyase | Sec/SPII |
| 000404 | PL6_1 |  | Versatile alginate lyase | Sec/SPI |
| 000406 | PL6 |  | Versatile alginate lyase | Sec/SPI |
| 000879 | PL38 |  | Endo-alginate lyase | (-) Globular |
| 001628 | PL17+CBM16/35/70 |  | Versatile alginate lyase | Sec/SPI |
| 001646 | PL17+CBM6 |  | Versatile alginate lyase | Sec/SPII |
| 002984 | PL7 |  | Versatile alginate lyase | Sec/SPI-SPII |
| 003205 | PL38 |  | Endo-alginate lyase | Sec/SPI-SPII |
| 003733 | PL17 |  | Versatile alginate lyase | Sec/SPII |
| 003800 | PL17+CBM16 |  | Versatile alginate lyase | Sec/SPI |
| 002469 | GH88 | Beta-1,4 uronidase | Exo-β-1,4-alginate oligomer hydrolase | (-) cytoplasmic |
| 004163 |  |  |  |  |
| Other polysaccharide lyases related to alginate lyases |  |  |  |  |
| 000048 | PL29 | Chondroitinase | Chondroitin sulfate ABC/hyaluronan | Sec/SPI |
| 000052 | PL29 | Chondroitinase | Chondroitin sulfate ABC/hyaluronan | Sec/SPI |
| 001723 | PL21 | Heparinase II | Heparin sulfate | Sec/SPI |
| 001725 | PL21 | Heparinase II | Heparin sulfate | Sec/SPI |
| 002230 | PL8_3 | Hyaluronidase | Hyaluronan | Sec/SPI |
| 002245 | PL35 | Chondroitinase | Chondroitin sulfate (in bacteria) | Sec/SPI |
| 002257 |  |  |  | (-) Globular |
| 002268 |  |  |  | (-) cytoplasmic |
| Transport of the alginate hydrolysis products |  |  |  |  |
| 001433-001438 | TonB/2 x TolQ/ExbB/TolR/ExbD/TonB plug | OM channel/OM transporter/ Energy transducer | TonB-dependent transport system: outer membrane transport of oligomers | Sec/SPI |
|  |  |  |  | Sec/SPIII membrane |
| 004062-004064 | TolQ/ExbB + TolR/ExbD/TonB plug | OM transporter/ Energy transducer |  |  |
| 004214-004215 | TolQ/ExbB + TolR/ExbD | OM channel OM transporter |  |  |
| 000051 | SglT/ToaA | Na<sup>+</sup> symporter SSF | Putative uronate transporter | CPM |
| Cytoplasmic uronate metabolism |  |  |  |  |
| 004145 | DehR | NADPH-dependent oxidoreductase | 4-deoxy-L-erythro-5-hexoseulose uronate>2-keto-3-deoxy-D-gluconate |  |
| 000255001079003637 | KduD/GndA | 2-keto-3-deoxygluconate oxidoreductase | DehR homologue |  |
| 000038003081003506 | KdgK | Sugar phosphokinase | 2-keto-3-deoxy-D-gluconate>2-keto-3-deoxy-6-phosphogluconate |  |
| 000037002471003082 | KdgA/KDGP aldolase | Sugar aldolase | 2-keto-3-deoxy-6-phosphogluconate>pyruvate + glycerol-3-phosphate |  |
| 002140 | KduI/UxaC | Uronate isomerase | glucuronate><fructuronate |  |
| 002468002470 | KdgF KdgR/GntR | Cupin Cupin+HTH-3 | putative pectin-utilization protein transcriptional regulator of the Kdg/Kdu pathway |  |
| 000254003636000517 | UxuA | Mannonate dehydratase | mannonate>2-keto-3-deoxy-D-gluconate |  |
| 000518000519 | UxuB UxaC/KduI | Mannonate dehydrogenase Uronate isomerase | mannonate>fructuronate glucuronate><fructuronate |  |

**Fig. 3.**
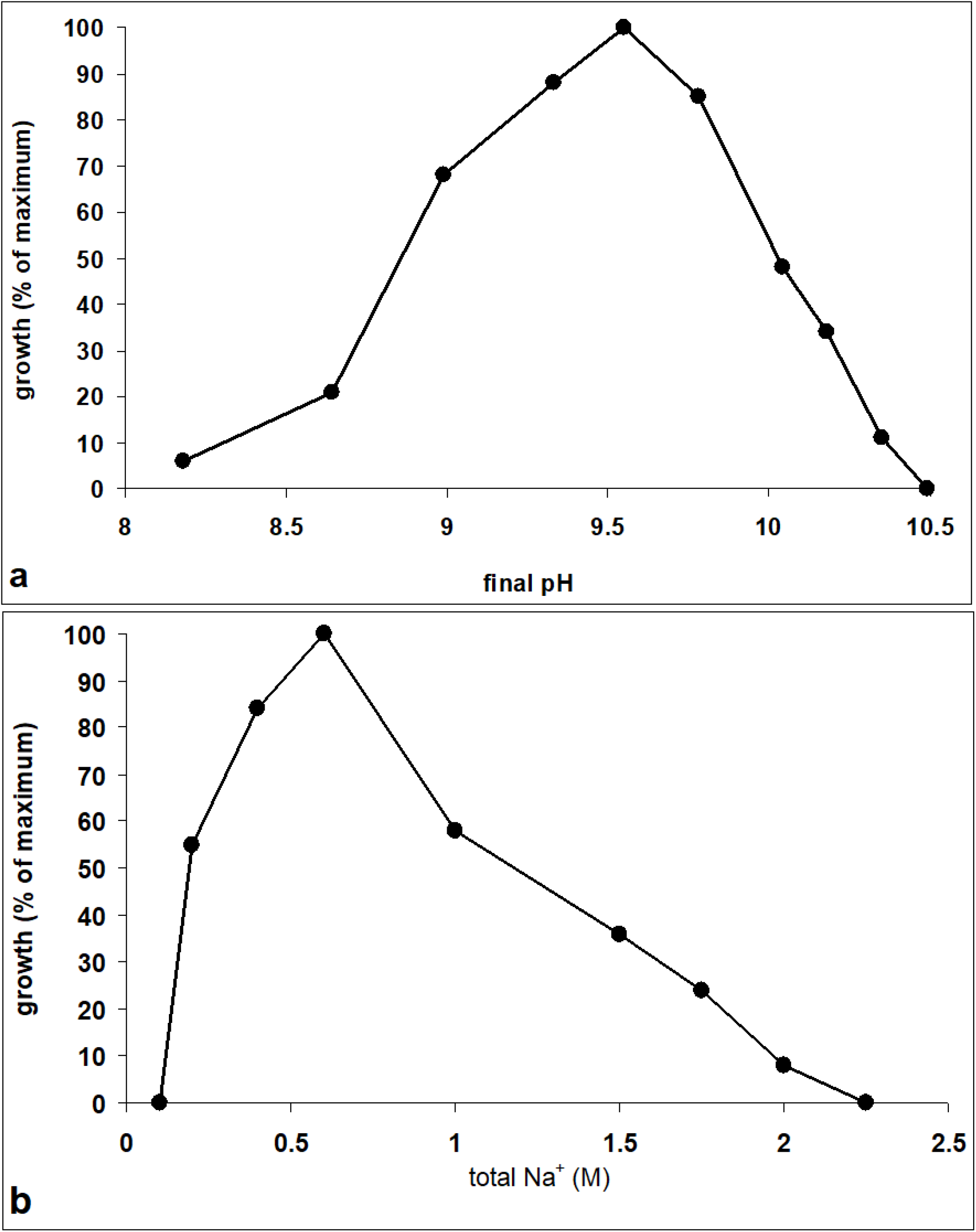
pH (at 0.6 M Na^+^) and salinity as sodium carbonates (at pH 9.5) for growth of strain AB-alg1^T^ with alginate at 30 ^°^C. The biomass was estimated by measuring OD_600_ (average values from a duplicate experiment).

### Functional genome analysis of strain AB-alg1^**T**^

#### Alginate utilization potential

For the well characterized alginate-utilizing bacteria among marine *Bacteroidota* and *Gammaproteobacteria*, initial depolymerization of alginate is achieved by the concert action of various extracellular polysaccharide lyases (PL) of endo- and exo-types currently classified in PL families 6, 7, 12, 14, 15, 17, 18, 31, 34, 38, 39, and 44 of the CAZy database. The resulting alginate oligomers and dimers are further converted into unsaturated monouronate by glycoside hydrolases from the GH88 family located either in the periplasm or cytoplasm (Hashimoto et al., 1999; He et al., 2022; Nishiyama et al., 2021; Xu et a l., 2021; Zhu and Yin, 2015).

dbCAN and HMMER searches supplemented with Uniprot Blast predicted potential alginate lyases family proteins encoded in the genome of AB-alg1^T^, including PL6, 7, 12, 14, 15, 38 and 39, most with a Sec/SPI signal peptide, indicating their extracytoplasmic location. (**Table 2**). Furthermore, a single GH88 family of unsatured uronyl oligohydrolase (apparently cytoplasmic) was identified, as well as a potential Na^+^:oligoalginate symporter “ToaA” from the SSF superfamily (He et al., 2022). The alginate lyase encoding genes in the AB-alg1^T^ genome were not assembled into a canonical polysaccharide utilization locus (PUL), as is often the case with the alginate-utilizing bacteria, but, still, several of them were paired as a (endo-) + (exo-) types.

The known pathway of alginate metabolism in Gammaproteobacteria consists of first importing alginate oligomers through the outer membrane via porins (TonB or SusCD systems (Blanvillain et al., 2007). Next, final hydrolysis of the oligomers into C6-uronate – 4-deoxy-L-erythro-5-hexoseulose uronic acid, DEHU – is performed by GH88 unsaturated uronyl hydrolase, either in the periplasm or cytoplasm. Finally, reductive cleavage of DEHU into pyruvate and 3-phosphoglycerate is performed by the Kdu-Kdg system (He et al., 2022; Nishiyama et al., 2021; Xu et al., 2021). Homologous proteins similar to all the enzymes involved in this pathway were identified in the AB-alg1^T^ genome, including a TonB-dependent outer membrane porin, GH88, DEHU reductase (DehR), 2-dehydro-3-deoxy-phosphouronate kinase (KdgK), and 2-keto-3-deoxy-6-phosphouronate aldolase (KdgA, part of the Entner-Doudoroff pathway). In some cases, multiple similar proteins were present in the genome. To test if the pathway was expressed in the presence of alginate, we grew strain AB-alg1^T^ with alginate, sucrose and mannose in parallel incubations, extracted and digested protein and analyzed expression using Orbitrap mass spectrometry. Many of the identified proteins were highly expressed and upregulated in the presence of alginate compared to growth on mannose and sucrose (**Table 2**).

The genome featured two clusters of genes potentially involved in alginate degradation that appeared to be upregulated as single operons in response to alginate: pgaptmp_000046-60 and pgaptmp_02465-70 (**Table S3**). The first appeared to be mainly focused on depolymerization and import, containing endo- and exo-PL families 15 and 39 alginate lyases, two PL29 chondroitinases, and a putative uronate transporter, among other genes (**Table 2**). The second appeared to be mainly focused on further degradation of the monomers in the cytoplasm and contained beta-1,4 uronidase, KdgA/KDGP aldolase/putative pectin-utilization protein, and a transcriptional regulator of the Kdg/Kdu pathway.

To explore the evolutionary history of these gene clusters, phylogenetic trees of genes from both upregulated regions were reconciled with the species tree using AleRax (Morel et al., 2024). Ancestral reconstruction (**Fig. 4**) showed that homologs of the alginate degradation genes from the gene cluster [pgaptmp_000047–000060] occur in all T3Sed10-336 MAGs from Kulunda soda lakes, but not in those from Canadian soda lakes (GCA_022565335, GCA_022565675, and GCA_020831535). It is possible that AB-alg1’s Canadian relatives feed on a different polysaccharide. Among other *Opitutales*, [pgaptmp_000047–000060] was only (nearly) complete in *Oceanipulchritudo coccoides*, a known alginate consumer (Feng et al., 2020), and *Cerasicoccaceae*. A broader search for polysaccharide lyase gene families among *Opitutaceae, Oceanipulchritudinaceae*, and T3Sed10-336 suggested that use of alginate and related acidic polysaccharides might be widespread among all three families (**Table S4**). So far, growth with alginate was not tested in any of the isolated species and only a single organism tested positive for alginolytic activity (*Oceanipultrichido*). Future research might investigate use of alginate more systematically for members of the *Opitutales*. The most-well represented alginate lyase type among the analyzed genomes was PL38, which is also widespread among marine *Verrucomicrobiota* (Lozada et al., 2025).

**Fig. 4.**
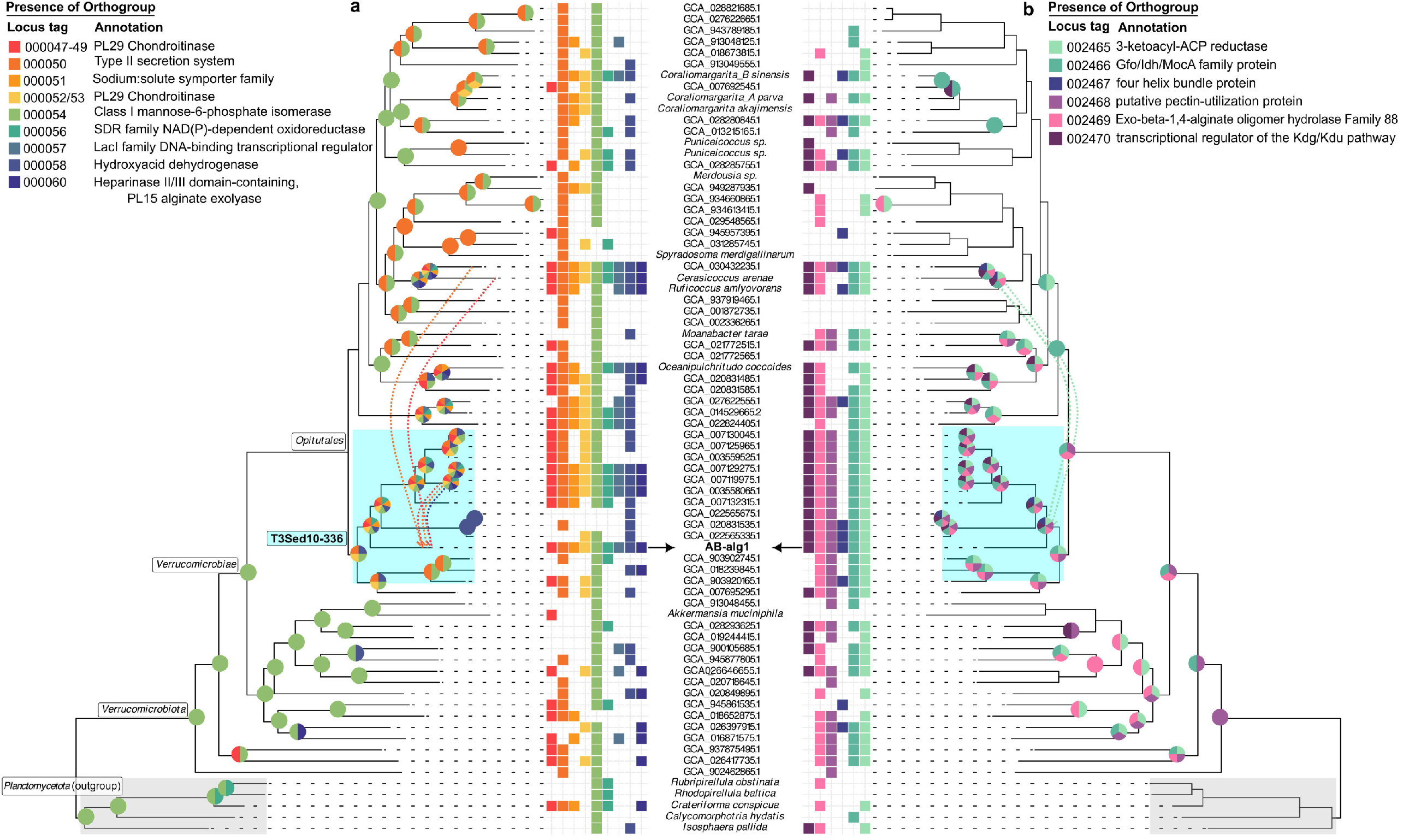
Reconciled species tree with presence of orthogroups associated with operons pgaptmp_000046-60, responsible for alginate depolymerization (**a**) and pgaptmp_02465-70, responsible for monomer degradation (**b**). Heatmaps show which orthogroups are present in the leaf genomes, while pie charts show which orthogroups are present at the ancestral nodes. Orthogroups associated with locus tags are given in **Table S5**. T3Sed10-336 is highlighted in cyan. Dashed arrows show horizontal gene transfer of [pgaptmp_000046-50, 60] from other T3Sed10-336 bacteria and [pgaptmp_000047–50, 2465] from *Cerasicoccaceae*. Conversely, ancestors of AB-alg1^T^ may have transferred [pgaptmp_002466] to *Cerasicoccaceae*. Note that *Cerasicoccaceae* have no current species adapted to alkaline environments.

AB-alg1^T^ had some unique features contributing to its alginolytic potential. First of all, the upregulated PL39 endoalginate lyase (pgaptmp_000055) did not have any close homologues among any of the genomes presented in **Fig. 4** (although distantly related, <50% identity, proteins were present occasionally). Second, AB-alg1^T^ obtained additional copies of pgaptmp_000047–000050 and pgaptmp_000060 from other T3Sed10-336 bacteria as well as pgaptmp_000047–000050 from *Cerasicoccaceae*. For the pulG-like type II secretion pseudopilin, pgaptmp_000050, no less than 67 homologous proteins were present in the AB-alg1^T^ genome, obtained in six gene-transfer events (**Fig. S1 and S2**), with remaining copies attributed to gene duplication. The type II secretion system (pgaptmp_004181-004193, upregulated the presence of alginate) could be involved in the export of the alginases pgaptmp_000055 and 000060, as was previously shown for cellulases in *Cytophaga hutchisonii* (Wang *et al*., 2017).

#### Other predicted key functional properties

The AB-alg1^T^ genome, as well as the genomes of related *Opitutales* (**Table S4**), encode multiple putative alginate lyases, and also other PL families with a potential hydrolytic activities against acidic glycosaminoglucan polysaccharides. Hyaluronate, chondroitin and heparin (PL 8, 29, 33, 35, 37) were most common. Furthermore, the AB-alg1^T^ genome encodes multiple copies (67) of desulfatases or sulfohydrolases, mostly clustered in three large loci. Although such enzymes are essential for depolymerization of the sulfated glucosaminoglucans chondroitin and heparin, our growth tests of both AB-alg isolates gave negative results with these polysaccharides. An attempt to use these three polysaccharides as selective enrichment substrates resulted in isolates from completely different phylogenetic lineages. The ability of AB-alg strains to use the β-fructan inulin is supported by the presence of two GH 32 inulinase genes (one excreted, one cytoplasmic) in the AB-alg1^T^ genome, while their growth on starch can be explained by multiple secreted GH13 amylases from subfamilies 10, 13, 14, 20, and 31.

As mentioned above, AB-alg1^T^ was able to grow by anaerobic respiration of nitrite (but not nitrate) to N_2_O with alginate as substrate. The genome encodes two types of dissimilatory nitrite reductase: a copper-containing denitrifying NirK (clade 5 in a bicistronic operon with its electron donor monoheme cytochrome *c*) and an ammonifying NrfAH. As we did not observe NH_3_ formation during anaerobic growth with nitrite, either with alginate or mannose as electron donors, the ammonifying pathway was not active, at least under the conditions tested. The denitrifying pathway from nitrite to N_2_O includes two enzymatic steps – nitrite reduction to NO by nitrite reductase (either Cu-containing NirK or cytochrome cd1 NirS) and NO reduction to N_2_O (either by quinol-dependent *q*NorB or cytochrome *c*-dependent (*c*NorBC). The AB-alg1^T^ genome encodes a NirK + its electron-accepting periplasmic cytochrome *c* and a homologue of *c*NorBC, belonging to the *e*Nor clade, according to the recent classification of Murali et al. (2025) (**Table 3**).

**Table 3.**
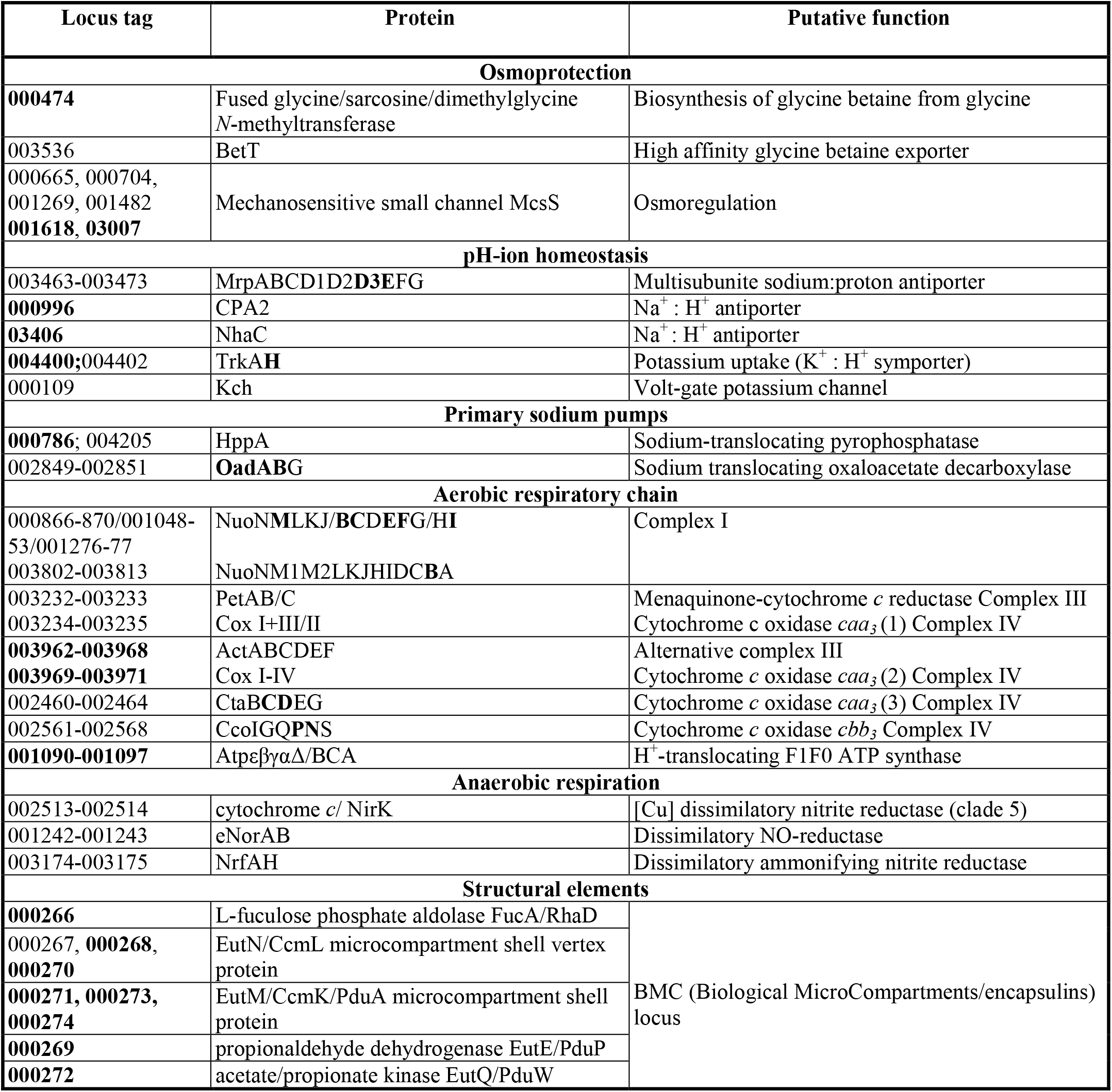
Proteins involved in haloalkaliphilic adaptation, energy metabolism and structural functions encoded in the genome of strain AB-alg1^T^. In **bold**: proteins expressed in cells growing aerobically with alginate. The complete data are provided in Supplementary Table S3.

#### Haloalkalphilic adaptation, energy generation and structural features

A search for genes encoding biosynthesis of osmolytes identified a single candidate gene encoding a fused version of the two SAM-dependent *N*-methyltransferases catalyzing step-wise N-methylation of glycine to glycine betaine (Nyyssölä et al., 2001). In addition, external glycine betaine could be imported by a high-affinity BCC transporter BetT. AB alg1^T^ may also use mechanical osmoregulation, based on the presence of six copies of genes coding for mechanosensitive small channel proteins McsS (**Table 3**). With respect to pH and ion homeostasis, the AB-alg1^T^ genome features the multisubunit Na^+^: H^+^ antiporter Mnh/MrpEFGBCD1D2D3, two single-subunit Na^+^: H^+^ antiporters CPA2 and NhaC, and a K^+^: H^+^ symporter TrkAH. Furthermore, genes for two primary sodium pumps, including 2 copies of a sodium-translocating pyrophosphatase HppA and a sodium-translocating oxaloacetate decarboxylase OadABG, were identified. The latter is often present in anaerobic alkaliphilic bacteria (Buckel, 2001).

A large genomic locus (ten genes) containing marker for biological microcompartments (BMCs, Kerfeld et al., 2018) or encapsulins was present in the genome of AB-alg1^T^ (**Table 3**) and is nearly identical in composition to one found the hyaluronan-utilizing planctomycete *Natronomicrosphaera* (Sorokin et al., 2025). Such protein-shells are formed to enclose toxic metabolic pathways in bacteria, in particular toxic aldehydes and alcohols during glycolytic degradation of rhamnose and fucose (Axen et al., 2014; Kennedy et al., 2021). Thin sectioning did not show the typical BMC-like inclusions in the two isolated soda lake bacteria, at least when the cells were growing on their respective acidic polysaccharides.

## Conclusion

Strain AB-alg1^T^ and AB-alg4 are the first pure culture representatives of natronophilic *Verrucomicrobiota*, isolated from soda habitats, and, to the best of our knowledge, also a first documented case of *Verrucomicrobiota* capable of growth on alginate. Nonetheless, the results of comparative genomic analysis showed that several *Opitutales* members do have alginolytic potential and that genes upregulated for alginate degradation in AB-alg1^T^ might have been transferred from other lineages. Future research should explore utilization of alginate as a growth substrate in culturable representatives of this phylum, especially those from marine habitats that provide for abundant alginate.

Based on distant phylogenomic and unique phenotypic properties, the natronophilic alginotrophic strains AB-alg1^T^ and AB-alg4 from soda lakes and soda solonchak soils, respectively, are proposed to be classified into a new genus and species *Verruconatronum alginivorum* gen, nov., sp. nov. within a new family *Verruconatronaceaea* fam. nov., which also incorporates several genus-level groups of MAGs from similar soda lake habitats, currently known as TS3-sed10 in GTDB. *Verruconatronaceae* forms a sister family to *Opitutaceae w*ithin the order *Opitutales*,

### Description of *Verruconatronaceae* fam. nov

[Ver.ru.co.na.tro.na’ceae. N.L. neut. n. *Verruconatronum*, the type genus of the family; -*aceae* ending to denote a family; N.L. fem. pl. n. *Verruconatronaceae*, the family of the genus *Verruconatronum*]

The family *Verruconatronaceae* includes facultatively anaerobic, moderately salt-tolerant and obligately alkaliphilic organoheterotrophic bacteria living in aquatic and terrestrial haloalkaline habitats. The members are saccharolytic, utilizing polysaccharides and simple sugars for aerobic or fermentative growth or anaerobic growth by denitrification. A particularly characteristic substrate is alginate from brown algae. The family includes a type genus *Verruconatronum* consisting of cultivated species and three genus-level groups comprised of MAGs from soda lakes in southwestern Siberia and Canada. The family is a member of the order *Opitutales*, class *Opitutia*, phylum *Verrucomicrobiota*.

### Description of *Verruconatronum* gen. nov

[Ver.ru.co.na.tro’num. L. fem. n. *verruca*, a wart; N.L. neut. n. *natron*, soda (arbitrarily derived from the Arabic n. *natrun* or *natron*); N.L. neut. n. *Verruconatronum*, soda-loving *Verrucomicrobiota* genus]

The genus includes moderately salt-tolerant, alkaliphilic, facultatively anaerobic, heterotrophic bacteria specialized on utilization of alginate as growth substrate. They are small motile cocci without pigmentation. Currently includes a single species based on two pure cultures isolated from soda lakes and soda solonchak soil. The genus is classified within its own family *Verruconatronaceae*, order *Opitutales*. The type species is *Verruconatronum alginivorum*.

### Description of *Verruconatronum alginivorum* sp. nov

[al.gi.ni.vo’rum. N.L. neut. n. *acidum alginicum*, alginic acid; N.L. neut. adj. suff. -*vorum*, devouring; from L. v. *voro*, to devour; N.L. neut. adj. *alginivorum*, alginic acid-devouring.

Cells are cocci, 0.7-1.0 μm, motile by 1-2 flagella with the Gram-negative type of the cell wall ultrastructure. The colonies are hard, white, up to 2 mm, forming inside of soft agar with concentration 0.6-0.8%. The polar membrane phospholipids are dominated by phosphatidylcholine and diphosphatidylglycerols (cardiolipins). Minor components include phosphatidylethanolamines, lyso-phosphatidylcholines, phosphatidylgylcerols and its acyl derivative. The dominant polar lipid fatty acids (in order of abundance) are *anteiso*15:0, 18:1ω9, 18:0 and 16:0. The only respiratory lipoquinone is MK-7:7. Has a genetic potential to synthesize osmolyte glycine betaine from glycine and to produce Bacterial MicroCompartments (BMC). Facultatively anaerobic saccharolytic bacteria growing best with alginate and mannose. Can also utilize starch, beta-fructan inulin and simple sugars, including glucose, fructose, fucose, sucrose, maltose, cellobiose and melezitose. Anaerobic fermentative growth was possible with alginate and mannose, and anaerobic respiration - with nitrite as the *e*-acceptor (N_2_O as the final product). Ammonium, urea and nitrate serve as the nitrogen source. Moderately salt-tolerant with the total Na^+^ (as sodium carbonates) range from 0.2 to 1.8 M of (optimum at 0.4-0.6 M). Obligate alkaliphilic with a pH range for growth from 8.2 to 10.3 (optimum at 9.5). Mesophilic, with the upper temperature limit for growth (at optimal pH and salinity) at 40 °C. The G + C content of the genomic DNA is 59.4 %. The type strain, AB-alg1^T^ (JCM 35393^T^=UQM 41574^T^), was isolated from a mix sample of surface sediments and brines from soda lakes in Kulunda Steppe (Altai, Russia) and an additional strain AB-alg4 (identical to the type strain in its 16S rRNA gene sequence) – from soda solonchak soils in the same area. The GenBank 16S-rRNA gene sequences of AB-alg1^T^ and AB-alg4 are PX129057 and PX129056, respectively. The genome size of the type strain is 6.01 Mb, the GenBank assembly number is XXXXXXX.

## Legends to the figures

**Fig. S1**. Ancestral reconstruction of pgaptmp_000046-60, showing transfer of several genes into the AB-alg1^T^ lineage. For each gene, colored line show co-speciation of genes and species.

**Fig. S2**. Ancestral reconstruction of pgaptmp_02465-70, showing transfer of pgaptmp_02467 into the AB-alg1^T^ lineage. For each gene, colored line show co-speciation of genes and species.

